# Can AI modelling of protein structures distinguish between sensor and helper NLR immune receptors?

**DOI:** 10.1101/2024.11.24.625045

**Authors:** AmirAli Toghani, Raoul Frijters, Tolga O. Bozkurt, Ryohei Terauchi, Sophien Kamoun, Yu Sugihara

## Abstract

NLR immune receptors can be functionally organized in genetically linked sensor-helper pairs. However, methods to categorize paired NLRs remain limited, primarily relying on the presence of non-canonical domains in some sensor NLRs. Here, we propose that the AI system AlphaFold 3 can classify paired NLR proteins into sensor or helper categories based on predicted structural characteristics. Helper NLRs showed higher AlphaFold 3 confidence scores than sensors when modelled in oligomeric configurations. Furthermore, funnel-shaped structures—essential for activating immune responses—were reliably predicted in helpers but not in sensors. Applying this method to uncharacterized NLR pairs from rice, we found that AlphaFold 3 can differentiate between putative sensors and helpers even when both proteins lack non-canonical domain annotations. These findings suggest that AlphaFold 3 offers a new approach to categorize NLRs and enhances our understanding of the functional configurations in plant immune systems, even in the absence of non-canonical domain annotations.

## INTRODUCTION

NLR (nucleotide binding and leucine-rich repeat) proteins are intracellular immune receptors that occur across all kingdoms of life but are particularly highly diversified in plants (Barragan and Weigel, 2021). These proteins function as singletons, pairs, or networks (Adachi et al., 2019b; Contreras et al., 2023a). Singleton NLRs can detect pathogens and execute hypersensitive cell death and immune responses, while paired NLRs have subfunctionalized into sensor (pathogen detection) and helper (immune execution) NLRs that carry distinct biochemical activities. In monocots, paired NLRs typically emerge from distinct phylogenetic clades to function together and do not have a common phylogenetic origin, unlike NLR networks (Contreras et al., 2023a; Kourelis et al., 2021). Sensor and helper NLR pairs are often genetically clustered, and some sensors have non-canonical integrated domains (IDs) that function in pathogen sensing and are absent in helper NLRs (Białas et al., 2018; Marchal et al., 2022). However, the presence of IDs is currently the only effective way to distinguish sensor NLRs from helper NLRs *in silico*, and no other method has been established to distinguish sensors from helpers in paired NLRs.

In plants, NLRs that carry a coiled-coil (CC) domain at their N-termini are the most widespread class. Following pathogen recognition, CC-NLR proteins oligomerize into pentameric or hexameric pore-like complexes (Förderer et al., 2022; Liu et al., 2024; Madhuprakash et al., 2024; Wang et al., 2019; Zhao et al., 2022). These complexes, known as resistosomes, are a defining feature of NLRs that execute the immune response such as the helpers; they translocate to cellular membranes and trigger immune responses such as calcium influx and hypersensitive cell death (Contreras et al., 2022; Duggan et al., 2021; Ibrahim et al., 2024). The prevailing model is that the funnel-shaped structure of CC-NLR resistosomes inserts into membranes and is required for cell death and immune response (Adachi et al., 2019b; Förderer and Kourelis, 2023; Wang et al., 2019). However, this funnel-shaped structure is formed by the N-terminal α1 helix of NLR, which is a structurally dynamic region that is difficult to resolve using cryo-electron microscopy (Förderer et al., 2022; Liu et al., 2024; Madhuprakash et al., 2024; Zhao et al., 2022). About 20% of plant CC-NLRs have a conserved sequence motif, called MADA, at the N-terminal α1 helix, while a previous study indicates that MADA motifs have degenerated in some sensor NLRs of solanaceous plants (Adachi et al., 2019a).

AlphaFold revolutionized protein structural biology by accurately modelling protein structures from primary sequences (Jumper et al., 2021). Recently, AlphaFold 3 (AF3) was released, enabling the modelling of protein structures with oleic acids serving as proxy for cellular membranes (Abramson et al., 2024). This function is particularly useful for proteins that dynamically interact with cellular membranes, such as CC-NLRs, and has enabled high-confidence modelling of CC-NLR resistosomes (Madhuprakash et al., 2024; Ibrahim et al., 2024). Here, we leveraged AF3 to explore the structural diversity of sensor and helper oligomers of a curated set of CC-NLRs consisting of i) experimentally validated NLR pairs in rice (Pikm, Pii, and Pia), ii) their orthologs (PIK5/6-NP, Pi5-3/1, and Pias), and iii) two previously cloned NLR pairs in barley (RPG5/HvRGA1 and RGH2/3) as summarized in **Table S1**. Our analyses revealed that helpers consistently exhibit higher AF3 confidence scores than sensors. Moreover, the funnel-shaped structures, crucial for the immune response, were consistently observed in helpers but not in sensors. We extended our investigations to uncharacterized NLR pairs of rice, and showed that AF3 can distinguish between NLR pairs even when both proteins do not carry any annotations for unconventional integrated domains. Based on these findings, we propose that AF3 can be used to classify paired NLRs into functional categories, providing a novel approach to understanding functional configurations of plant immune receptors.

## RESULTS

### Helper NLRs produce higher AlphaFold 3 pTM scores than their paired sensors

To benchmark the AF3 capacity to distinguish between sensor and helper NLRs, we curated eight NLR pairs in which the sensor can be readily assigned because it carries an ID (**Fig. S1**; **Table S1**). As in previous studies (Madhuprakash et al., 2024; Ibrahim et al., 2024), we used the oligomerizing domains of the NLR proteins, from the N-terminus to the end of the NB-ARC domain, and performed AF3 predictions of 5x and 6x stoichiometries with 50 oleic acids and using three different seed values. We then compared the sensor and helper predicted template modelling (pTM) scores (**Fig. 1a and b**; **Fig. S2–4**; **Table S2**). Helper NLRs consistently exhibited higher pTM scores than sensor NLRs in both pentameric and hexameric configurations, indicating that AF3 pTM scores can be used to distinguish between sensor and helper NLRs.

**Fig. 1.**
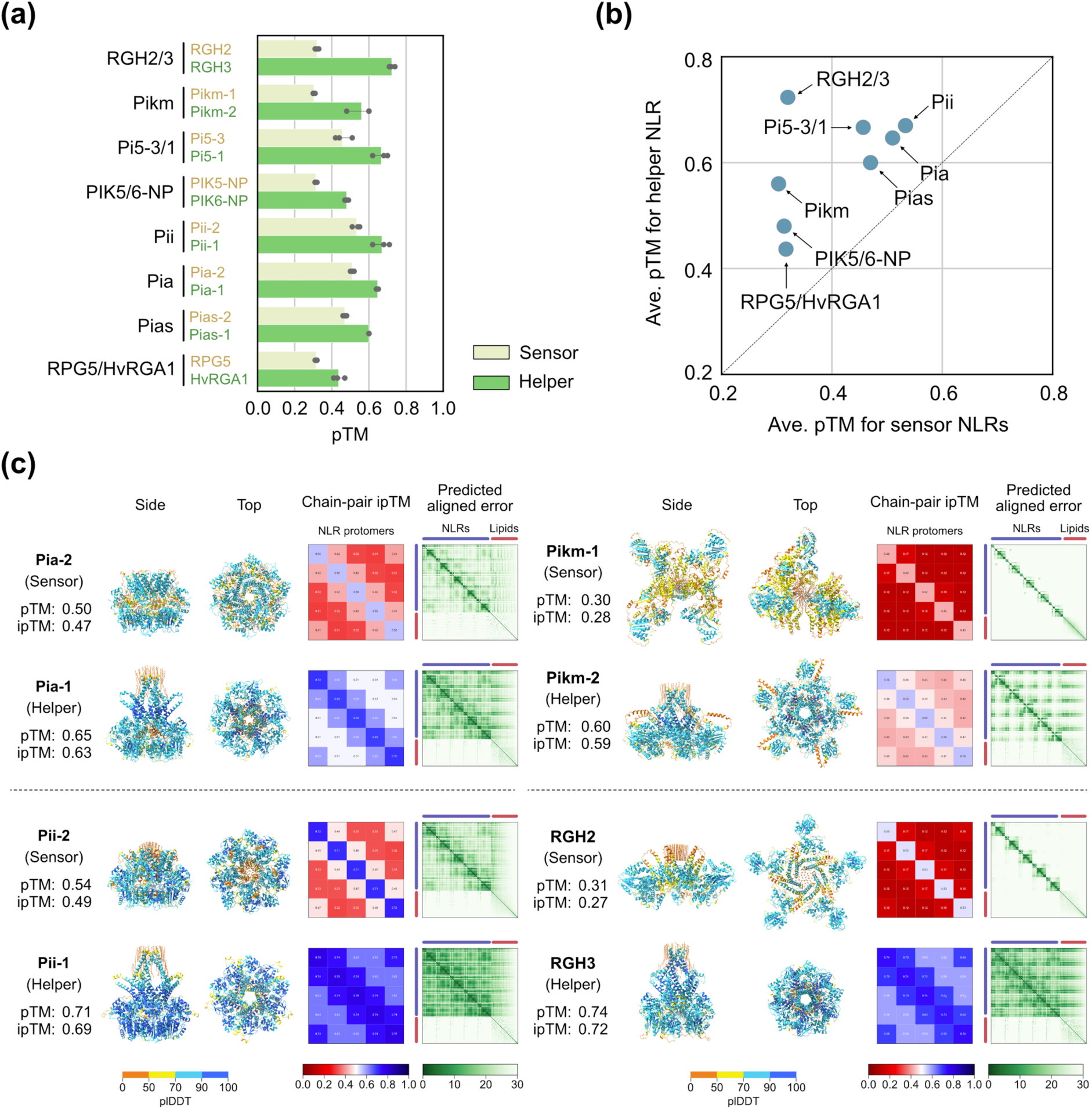
Helper NLRs produce higher AlphaFold 3 pTM scores than their paired sensors. a) Bar plot comparing sensor and helper pTM scores in pentameric AlphaFold 3 predictions. The amino acid sequences of the oligomerizing domains of the NLR proteins, from the N-terminus to the end of the NB-ARC domain, were used for the prediction. The pentameric structures were modelled with 50 oleic acids using three different seed values. b) Scatter plot comparing sensor and helper pTM scores in pentameric AlphaFold 3 predictions. The resulting pTM scores were averaged across three seed values for each sensor and helper NLR. c) Pentameric AlphaFold 3 structures of the four representative NLR pairs. The structures predicted using seed value 1 were visualized using ChimeraX (Meng et al., 2023).

### Helper AF3 structures form funnel-shaped structures unlike sensors

In addition to the AF3 confidence scores, we also examined the AF3 structures for distinct structural patterns between sensor and helper NLRs (**Fig. 1c**; **Fig. S5 and S6**). Helper NLRs consistently formed funnel-shaped structures in contrast to sensor NLRs. This observation aligns with previous studies demonstrating that sensor NLRs cannot execute the immune response on their own and may have lost the capacity to oligomerise into resistosome-like structures (Adachi et al., 2019b; Contreras et al., 2022). Sensors formed resistosome-like structures but with low confidence scores (**Fig. 1c**; **Fig. S5 and S6**). These findings further indicate that AF3 can distinguish between sensor and helper NLRs.

We then analyzed whether the presence of the MADA motif could classify sensors or helpers in the curated NLR pairs using the MADA HMM model in Adachi et al., 2019a (**Table S3**). The helpers of Pikm and PIK5/6-NP contained the MADA motif with HMM scores exceeding the cut-off value of 10, while the other sensors and helpers did not. Based on these results, only Pikm and PIK5/6-NP could be classified according to the presence of the MADA motif, whereas the remaining NLR pairs could not be classified. These findings indicate that AF3 modelling-based classification is more reliable than the sequence-based classification and highlight the utility of AF3 for NLR classification.

### AF3 can classify paired NLRs even when the putative sensors lack an integrated domain annotation

Based on Stein et al., 2018, we extracted ten rice CC-NLR pairs that i) are genetically linked in head-to-head orientations; ii) belong to distinct phylogenetic clades; and iii) carry a full N-terminal CC domain (**Fig. 2a**; **Fig. S1; Table S4–S6**). In five of the ten pairs, one of the NLRs carries an ID annotation and is presumed to be the sensor. The putative helpers (without ID annotation) had higher AF3 pTM scores than sensors (with ID annotation) for these pairs (**Fig. 2b**; **Fig. S7; Table S7**). Furthermore, putative helpers formed funnel-shaped structures, whereas putative sensors did not (**Fig. 2c**; **Fig. S8 and S9**). These results further confirm the application of AF3 for functional classification as noted with the eight previously characterized pairs.

**Fig. 2.**
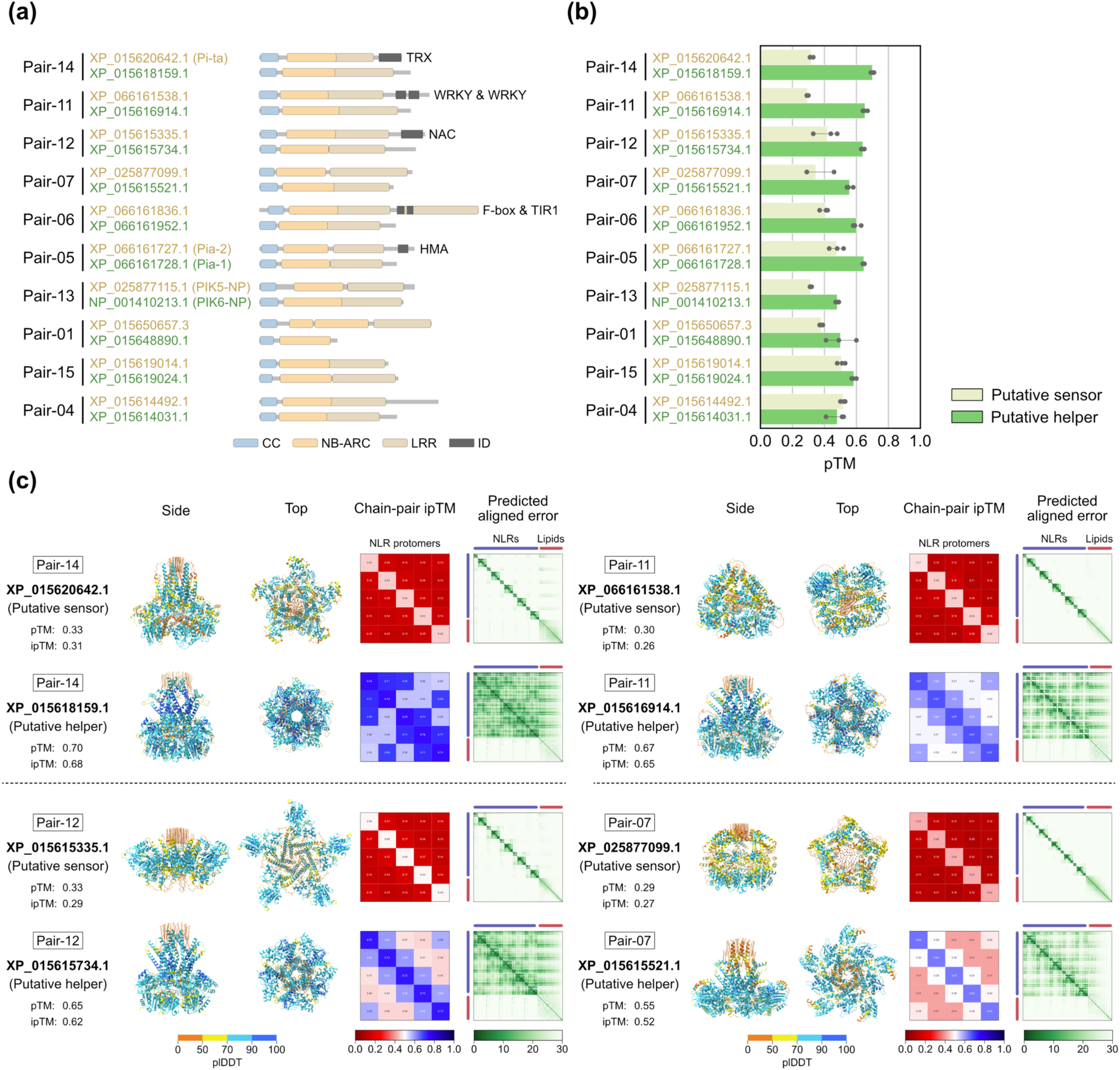
AF3 can classify paired NLRs even when the putative sensors do not carry an integrated domain annotation. a) Domain architectures of ten rice NLR pairs described by Stein et al., 2018. The domains were annotated with NLRtracker (Kourelis et al., 2021) and visualized with refplantnlR (https://github.com/JKourelis/refplantnlR). b) Bar plot comparing putative sensor and helper pTM scores in pentameric AlphaFold 3 predictions. The amino acid sequences of the oligomerizing domains of the NLR proteins, from the N-terminus to the end of the NB-ARC domain, were used for the prediction. The pentameric structures were modelled with 50 oleic acids using three different seed values. Putative sensors and helpers were assigned based on the average pTM scores of pentameric and hexameric structures, with a putative sensor having the lower average score and a putative helper having the higher average score. c) Pentameric AlphaFold 3 structures of the four representative NLR pairs described by Stein et al., 2018. The structures predicted using seed value 1 were visualized using ChimeraX (Meng et al., 2023).

Next, we examined the five remaining pairs whose NLRs do not carry any ID annotations. In each pair, we averaged the pTM scores of pentameric and hexameric configurations and assigned them as putative helpers or sensors based on higher or lower pTM scores, respectively (**Fig. 2b**; **Fig. S1 and S7; Table S7**). Pair-13 is identical to the rice pair PIK5/6-NP, in which PIK5-NP is known as a sensor with an integrated domain that is not annotated by InterProScan (Białas et al., 2021; Kourelis et al., 2021). Nonetheless, AF3 successfully classified them into helper or sensor based on the pTM scores even without the ID annotation. Among the other four pairs, Pair-07 exhibited a consistent difference in the pTM scores between the putative sensor and helper NLRs (**Fig. S7; Table S7**).

## DISCUSSION

AlphaFold 3 can potentially distinguish between sensor and helper NLRs based on their predicted structures and confidence scores. In AF3 predictions, helpers consistently exhibited higher AF3 confidence scores than sensors (**Fig. 1**; **Fig. S2–4**). Moreover, the funnel-shaped structures were consistently observed in helpers but not in sensors (**Fig. 1c and 2c; Fig. S5, S6, S8, and S9**). These findings indicate that AF3 might be able to distinguish the functional configurations of plant immune receptors. Our results highlight the utility of AF3 as a powerful tool for distinguishing structural and functional differences in paired NLRs.

Why did sensor NLRs exhibit lower AF3 confidence scores than helper NLRs? Previous studies proposed a model suggesting that paired NLRs evolved from singleton NLRs (Adachi et al., 2019b). In this model, singleton NLRs, which can detect pathogens (sensor) and execute hypersensitive cell death (helper), subfunctionalized into sensor or helper NLRs (Adachi et al., 2019b; Contreras et al., 2023a). This subfunctionalization results in the loss of cell death activity in sensor NLRs and the loss of pathogen perception activity in helper NLRs. Since the formation of resistosomes with funnel-shaped structures is essential for induction of cell death and other immune responses, the presence of stable resistosome and funnel-shaped structures in helper NLRs, but not in sensor NLRs, supports the previously proposed model. Consequently, sensor NLRs exhibit lower AF3 confidence scores than helper NLRs, reflecting their evolutionary divergence and functional specialization.

Recently, several researchers have applied AlphaFold to plant-pathogen interactions (Contreras et al., 2023b; Cruz et al., 2024; Ibrahim et al., 2024, 2023; Madhuprakash et al., 2024; Selvaraj et al., 2024; Seong et al., 2024; Sugihara et al., 2023; Tamborski et al., 2023). These advancements provide a novel approach to understanding functional configurations of immune receptors beyond domain architectures, sequence motifs, and phylogenetic relationships.

## METHODS

### NLR annotation

NLRs were annotated using NLRtracker v1.0.3 (Kourelis et al., 2021) and InterProScan v5.67-99.0 (Jones et al., 2014). Note that InterProScan did not annotate an integrated domain of PIK5-NP as previously reported (Kourelis et al., 2021). Therefore, we manually replaced the domain architecture of PIK5-NP from “CNL” to “CONL” as it contains the HMA domain between the NB-ARC and LRR (Białas et al., 2021). The domain architectures were visualized using refplantnlR (https://github.com/JKourelis/refplantnlR). The MADA motifs were analyzed using the HMM model in Adachi et al., 2019a and HMMER v3.4 (http://hmmer.org) with the option “--max”. The NLRtracker and HMMER outputs are archived on Zenodo (https://doi.org/10.5281/zenodo.13826775).

### Curation of NLR sequences

The sequences of curated NLR pairs were derived from either RefPlantNLR (Kourelis et al., 2021) or NCBI (**Table S1**). Regarding the NLR pairs described by Stein et al., 2018, the protein sequences of *Oryza sativa* cv. Nipponbare (Oryza_sativa_vg_japonica.protein.fasta) were downloaded from the URL (https://doi.org/10.7946/P2FC9Z). Based on Supplementary Data 6 in Stein et al., 2018, the NLR pairs that i) are genetically linked in head-to-head orientations; ii) belong to distinct phylogenetic clades were extracted (**Table S4**). Using DIAMOND BLASTP v2.1.9 (Buchfink et al., 2021), the corresponding NLR sequences were identified from the NCBI RefSeq annotation of rice cultivar Nipponbare genome (GCF_034140825.1). The best-hit sequences were summarized in **Table S5** and confirmed to be genetically linked in a head-to-head orientation. Based on NLRtracker outputs, the CC-NLR pairs, which carry a full N-terminal CC domain, were retained (**Table S6**). Piar-05, Piar-13, and the sensor of Pair-14 correspond to Pia, PIK5/6-NP, and Pi-ta, respectively (**Table S4 and S6**).

### AF3 prediction

Based on NLRtracker outputs, the amino acid sequences from the N-terminus to the end of the NB-ARC domain were extracted. Using the AlphaFold 3 webserver (https://golgi.sandbox.google.com), the extracted sequences were modelled in both pentameric and hexameric configurations with 50 oleic acids using three different seed values. The input sequences and resulting models are archived on Zenodo (https://doi.org/10.5281/zenodo.13826775).

### Phylogenetic analysis

We built the tree of monocot paired NLRs with RefPlantNLR (Kourelis et al., 2021) and a set of 4,936 NLR proteins from the NLRtracker output of 13 RefSeq proteomes, including two dicot species (*Arabidopsis thaliana* and *Solanum lycopersicum*) and 11 monocot species (*Zea mays*, *Triticum aestivum*, *Setaria viridis*, *Phragmites australis*, *Phoenix dactylifera*, *Oryza sativa*, *Musa acuminata*, *Lolium perenne*, *Hordeum vulgare* subsp. *vulgare*, *Brachypodium distachyon*, and *Asparagus officinalis*) (**Data S1, S2, S3, and S4**) (Toghani and Kamoun, 2024). More details on the phylogenetics analysis are available on GitHub (https://github.com/amiralito/Paired_NLR_AF3).

## ACKNOWLEDGEMENTS

We thank Andres Posbeyikian (The Sainsbury Laboratory) and Kentato Fujita (Osaka University) for valuable comments and scientific discussions.

## DATA AVAILABILITY

The scripts and dataset used for the phylogenetic analysis are available on GitHub (https://github.com/amiralito/Paired_NLR_AF3). The other datasets are archived on Zenodo (https://doi.org/10.5281/zenodo.13826775).

---

**Fig. S1.**
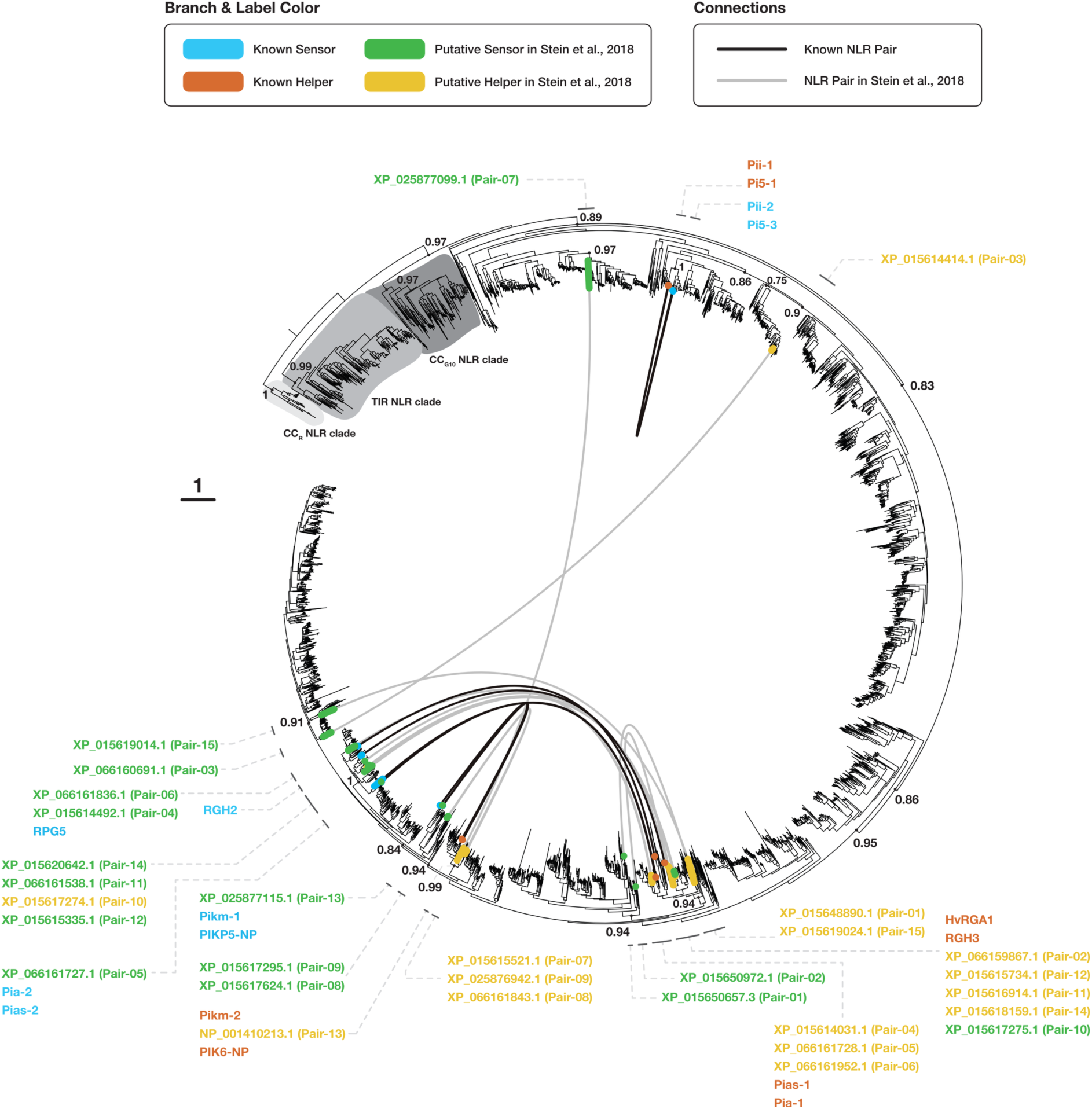
Phylogenetic tree of monocot paired NLRs with RefPlantNLR and 4,936 NLR proteins from 13 RefSeq proteomes, including two dicot species and 11 monocot species obtained from Toghani et al., 2024. The tree was built using the NB-ARC domains of the eight known NLR pairs summarized in **Table S1**, RefPlantNLR, and 4,936 NLR proteins from the NLRtracker output of 13 RefSeq proteomes, including two dicot species (*Arabidopsis thaliana* and *Solanum lycopersicum*) and 11 monocot species (*Zea mays*, *Triticum aestivum*, *Setaria viridis*, *Phragmites australis*, *Phoenix dactylifera*, *Oryza sativa*, *Musa acuminata*, *Lolium perenne*, *Hordeum vulgare* subsp. *vulgare*, *Brachypodium distachyon*, and *Asparagus officinalis*) (**Data S1, S2, S3, and S4**) (Toghani and Kamoun, 2024). The paired NLRs of rice cultivar Nipponbare (GCF_034140825.1), described by Stein et al., 2018, are included among 13 RefSeq proteomes. The genetic linkage between NLR pairs is indicated by a connection between linked nodes. Known sensor and helper NLR nodes are colored blue and orange, respectively, while putative sensor and helper nodes are colored green and yellow. Numbers next to the tree nodes indicate bootstrap values. The tree is rooted at the CC_R_-NLR clade. AlphaFold 3 was not performed for Pair-02, Pair-03, Pair-08, Pair-09, and Pair-10 because they do not carry a full N-terminal CC domain.

**Fig. S2.**
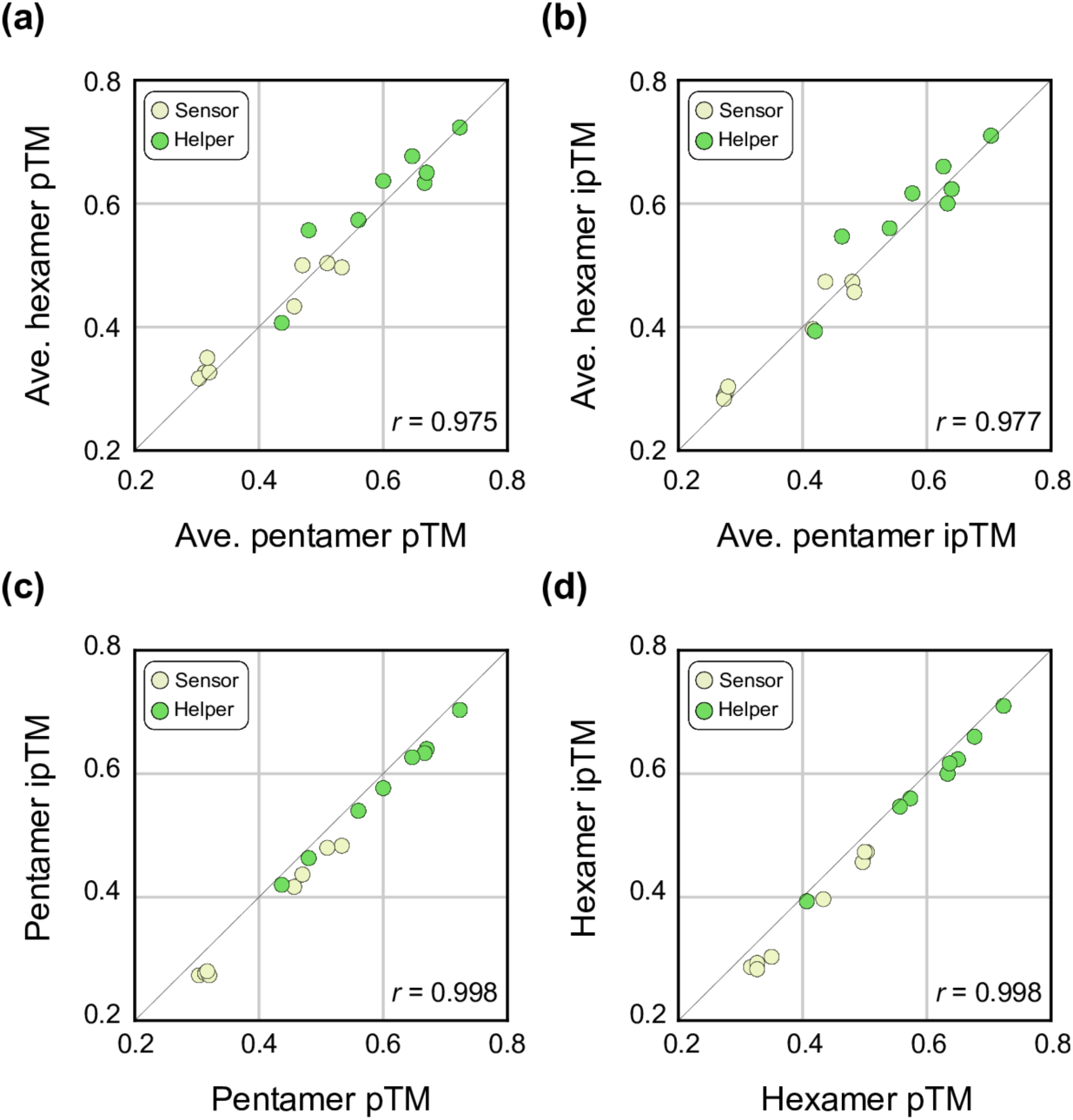
Correlations between pentameric and hexameric AlphaFold 3 confidence scores and between pTM and ipTM scores for previously reported NLR pairs. a) Correlation between the average pentameric and hexameric pTM scores. b) Correlation between the average pentameric and hexameric ipTM scores. c) Correlation between the pentameric pTM and ipTM scores. d) Correlation between the hexameric pTM and ipTM scores. The amino acid sequences of the oligomerizing domains of the NLR proteins, from the N-terminus to the end of the NB-ARC domain, were used for the prediction. The pentameric and hexameric structures were modelled with 50 oleic acids using three different seed values. The resulting pTM or ipTM scores were averaged across three seed values for each NLR in Fig. S2a and S2b. The Pearson rank correlation coefficients (*r*) were calculated with the SciPy library in Python.

**Fig. S3.**
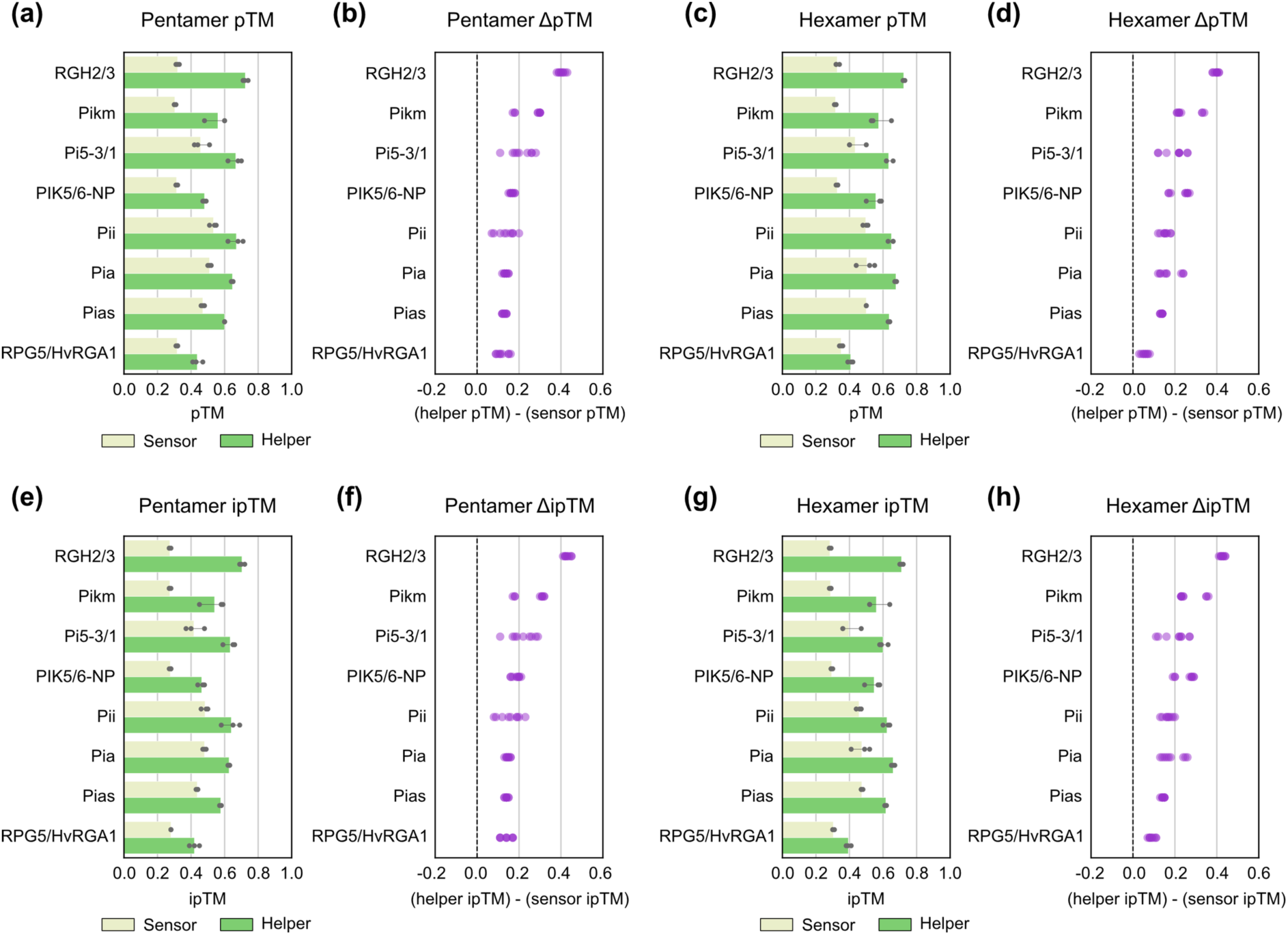
Comparisons of sensor and helper AlphaFold 3 scores in pentameric and hexameric configurations for previously reported NLR pairs. a-d) pTM score comparisons for pentamers (a,b) and hexamers (c,d). e-h) ipTM score comparisons for pentamers (e,f) and hexamers (g,h). The a, c, e, and g panels show bar plots, and the b, d, f, and h panels display score differences (helper minus sensor). The amino acid sequences of the oligomerizing domains of the NLR proteins, from the N-terminus to the end of the NB-ARC domain, were used for the prediction. The pentameric and hexameric structures were modelled with 50 oleic acids using three different seed values. Subtractions were performed for all possible pairs of sensor and helper scores.

**Fig. S4.**
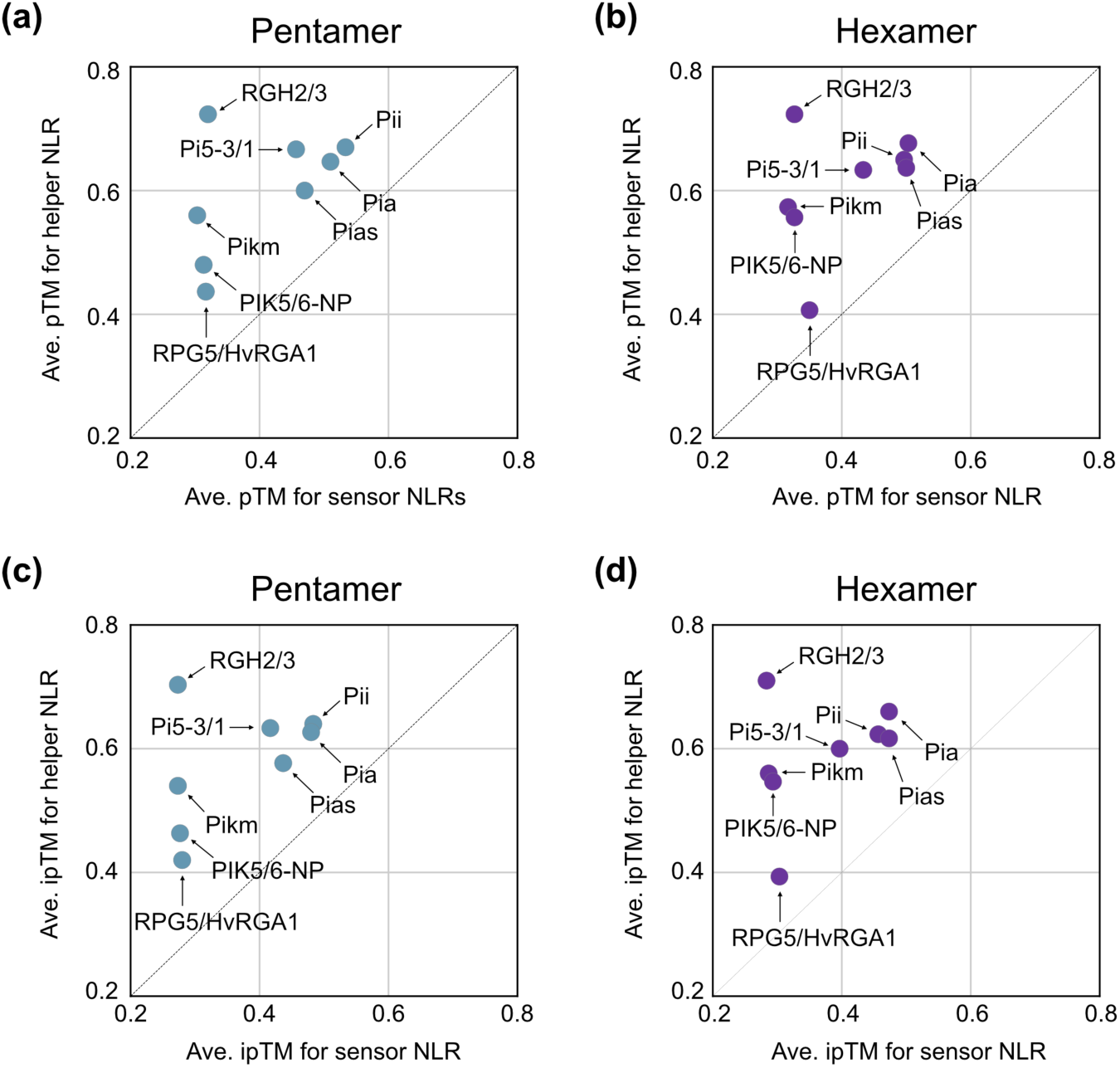
Scatter plots comparing average sensor and helper scores in AlphaFold 3 predictions. a) Scatter plot comparing average sensor and helper pTM scores in pentamer predictions. b) Scatter plot comparing average sensor and helper pTM scores in hexamer predictions. c) Scatter plot comparing average sensor and helper ipTM scores in pentamer predictions. d) Scatter plot comparing average sensor and helper ipTM scores in hexamer predictions. The amino acid sequences of the oligomerizing domains of the NLR proteins, from the N-terminus to the end of the NB-ARC domain, were used for the prediction. The pentameric and hexameric structures were modelled with 50 oleic acids using three different seed values. The resulting pTM or ipTM scores were averaged across three seed values for each sensor and helper NLR.

**Fig. S5.**
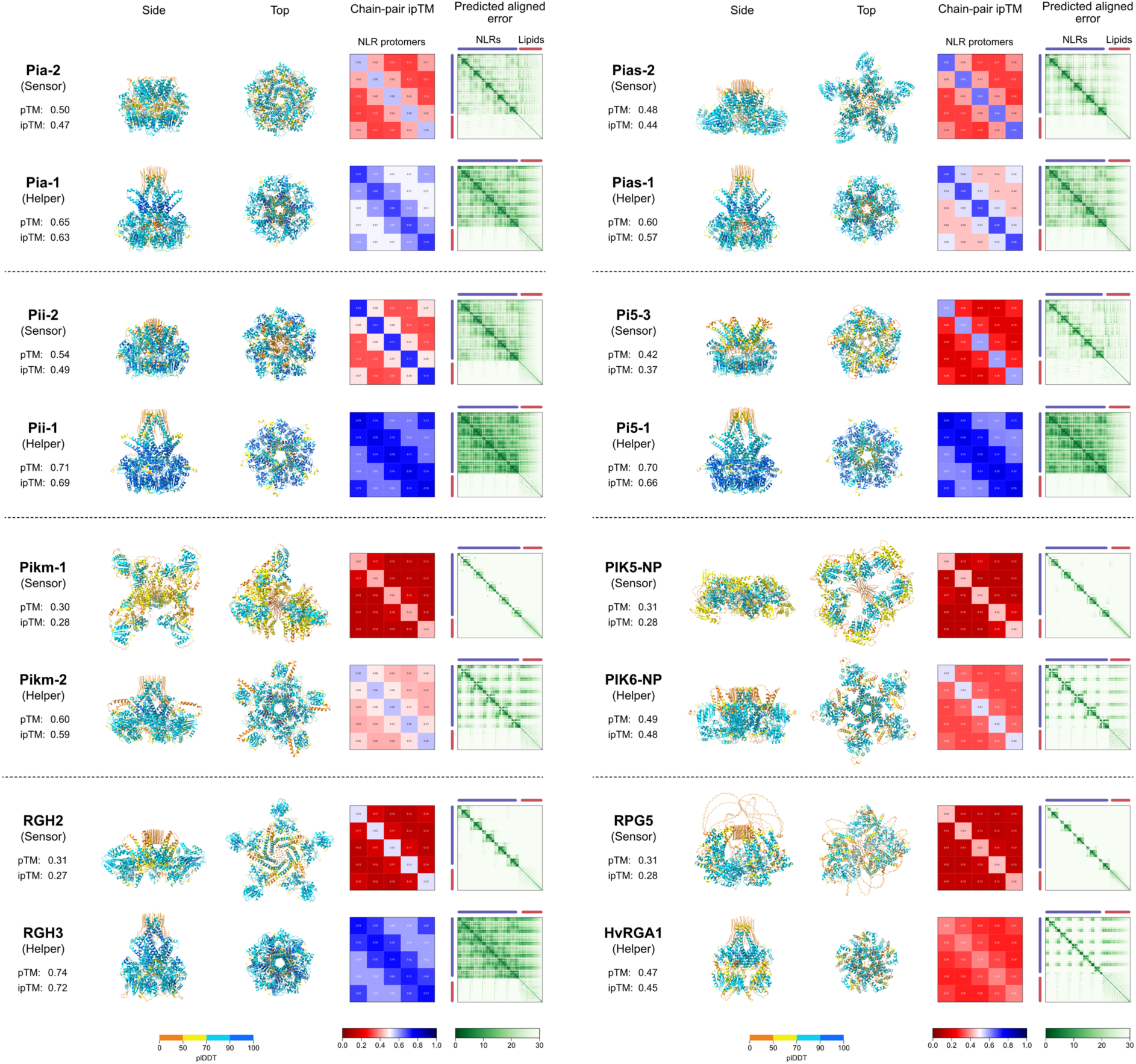
Pentameric AlphaFold 3 predictions for previously reported NLR pairs. The amino acid sequences of the oligomerizing domains of the NLR proteins, from the N-terminus to the end of the NB-ARC domain, were used for the prediction. The pentameric structures were modelled with 50 oleic acids using seed value 1. The predicted structures were visualized using ChimeraX (Meng et al., 2023).

**Fig. S6.**
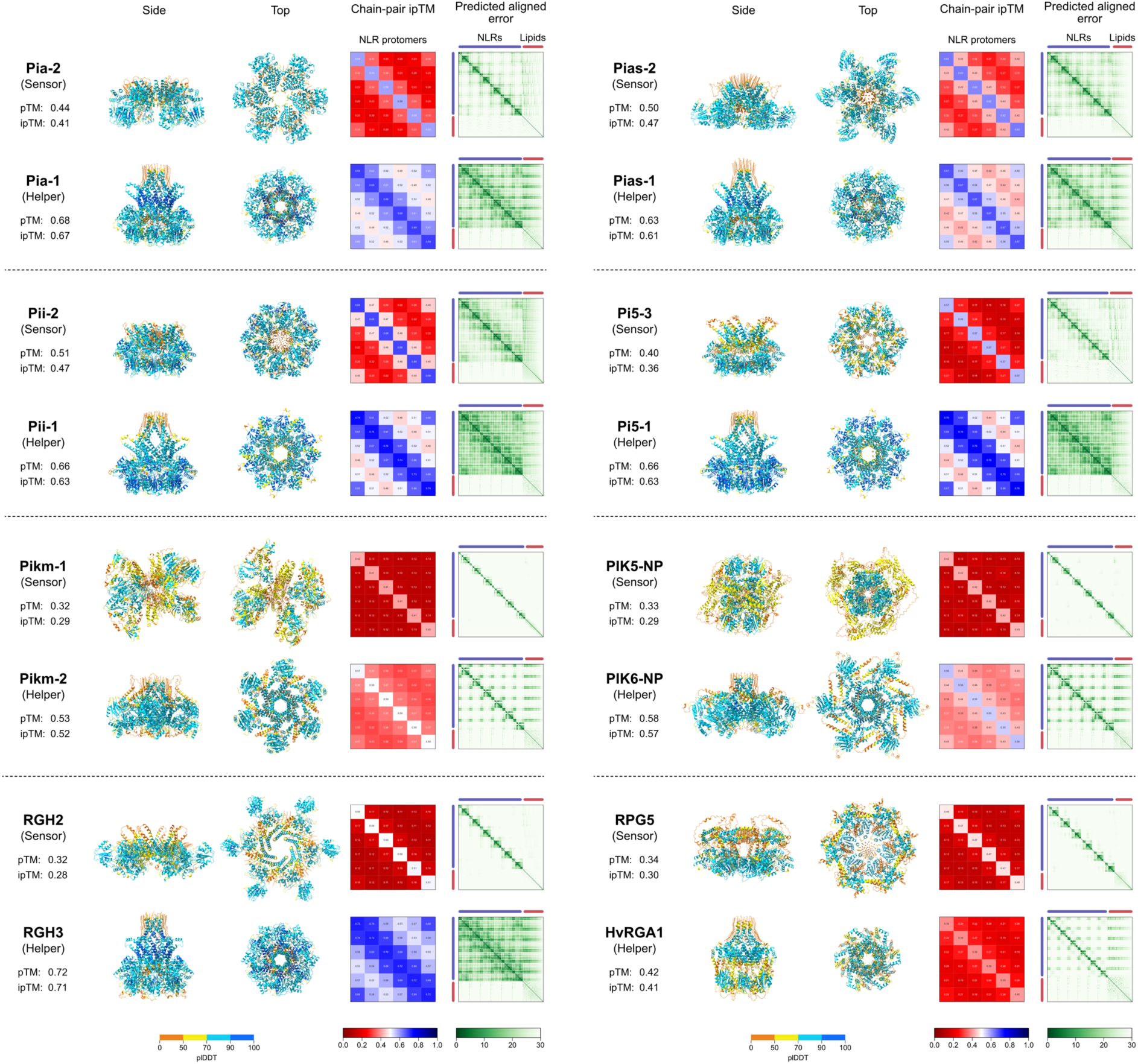
Hexameric AlphaFold 3 predictions for previously reported NLR pairs. The amino acid sequences of the oligomerizing domains of the NLR proteins, from the N-terminus to the end of the NB-ARC domain, were used for the prediction. The hexameric structures were modelled with 50 oleic acids using seed value 1. The predicted structures were visualized using ChimeraX (Meng et al., 2023).

**Fig. S7.**
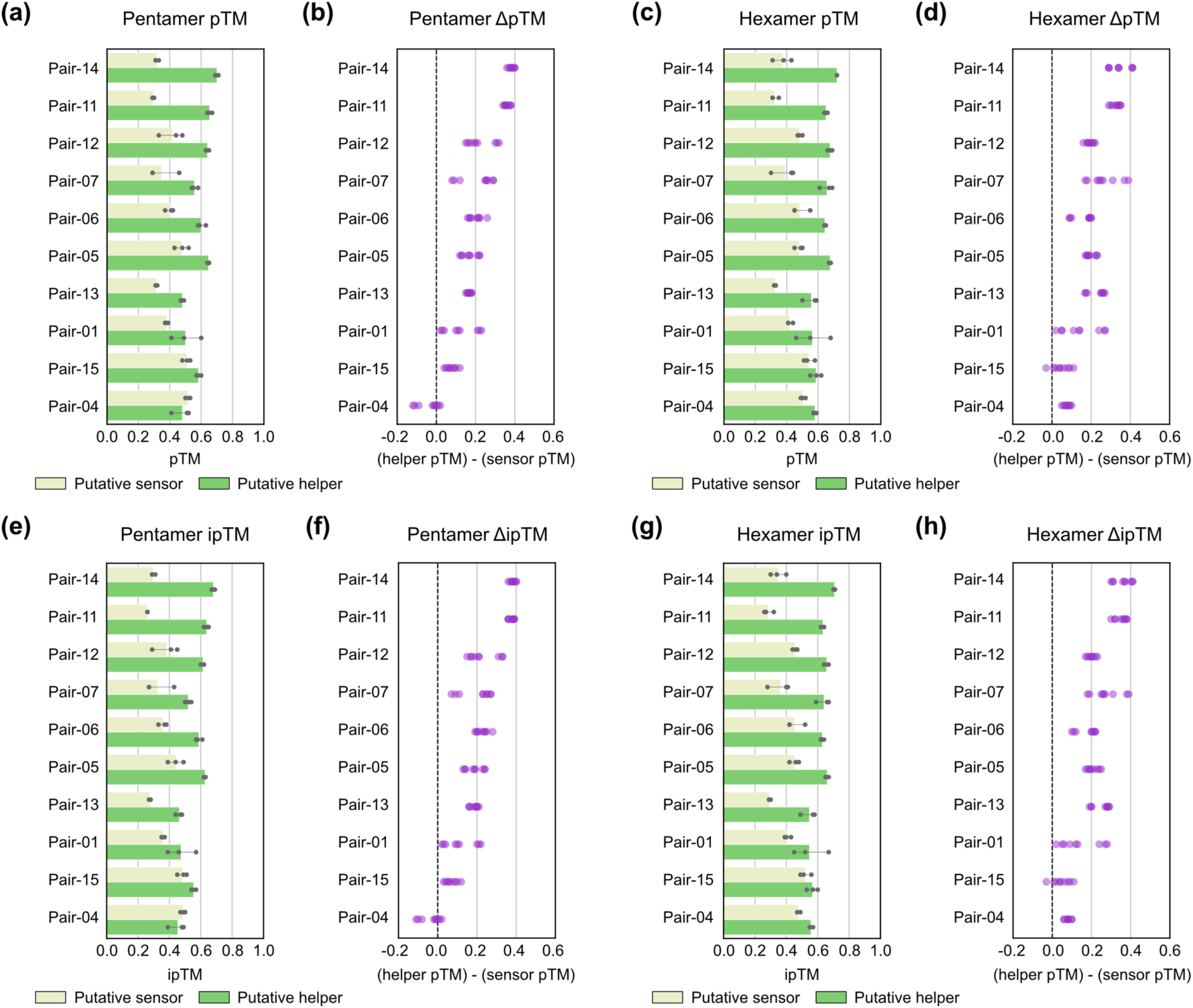
Comparisons of putative sensor and helper AlphaFold 3 scores in pentameric and hexameric configurations for NLR pairs described by Stein et al., 2018. a-d) pTM score comparisons for pentamers (a,b) and hexamers (c,d). e-h) ipTM score comparisons for pentamers (e,f) and hexamers (g,h). The a, c, e, and g panels show bar plots, and the b, d, f, and h panels display score differences (putative helper minus putative sensor). The pentameric and hexameric structures were modelled with 50 oleic acids using three different seed values. Putative sensors and helpers were assigned based on the average pTM scores of pentameric and hexameric structures, with a putative sensor having the lower average score and a putative helper having the higher average score. Subtractions were performed for all possible pairs of putative sensor and helper scores.

**Fig. S8.**
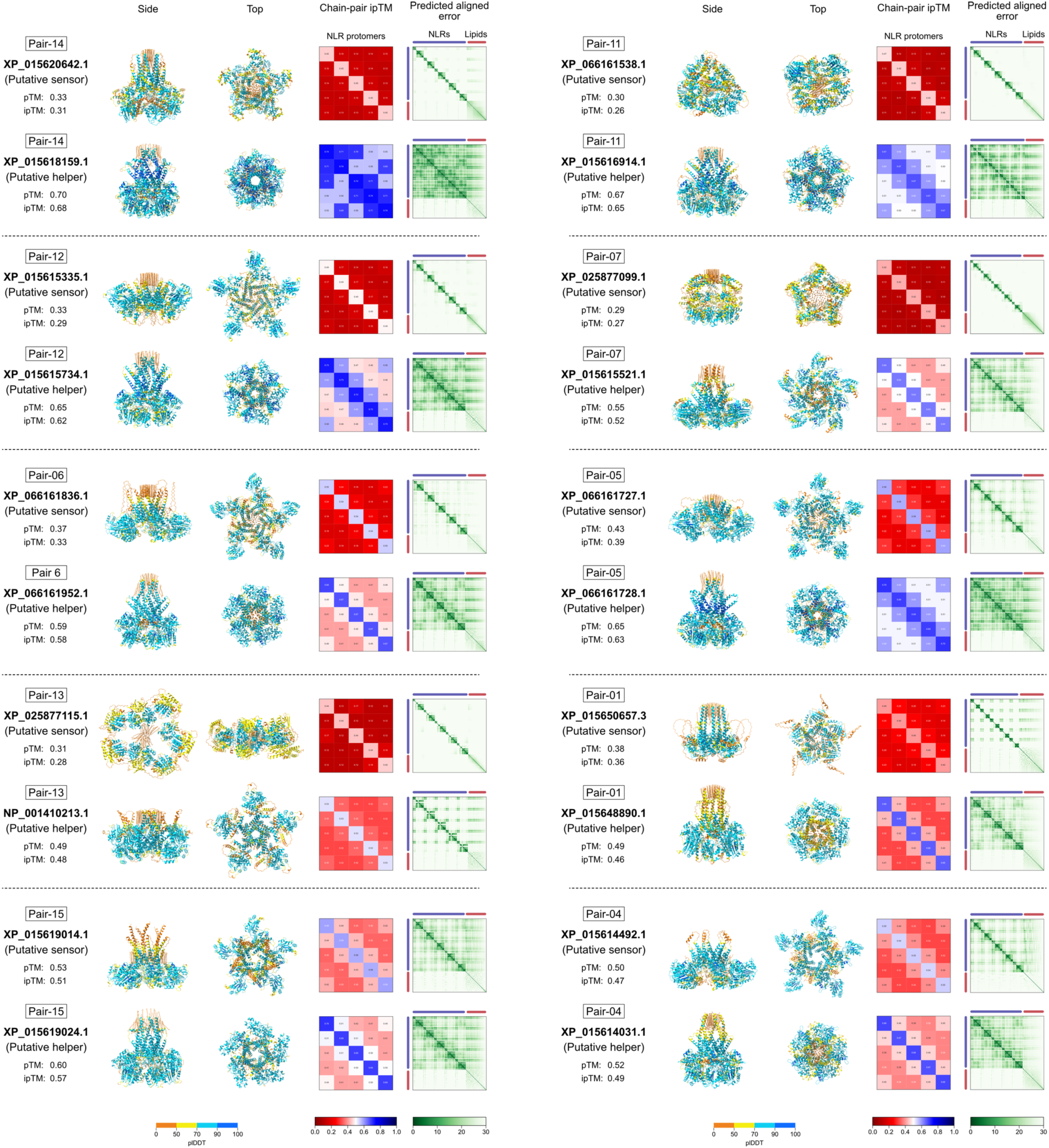
Pentameric AlphaFold 3 predictions for NLR pairs described by Stein et al., 2018. The amino acid sequences of the oligomerizing domains of the NLR proteins, from the N-terminus to the end of the NB-ARC domain, were used for the prediction. The pentameric structures were modelled with 50 oleic acids using seed value 1. The predicted structures were visualized using ChimeraX (Meng et al., 2023).

**Fig. S9.**
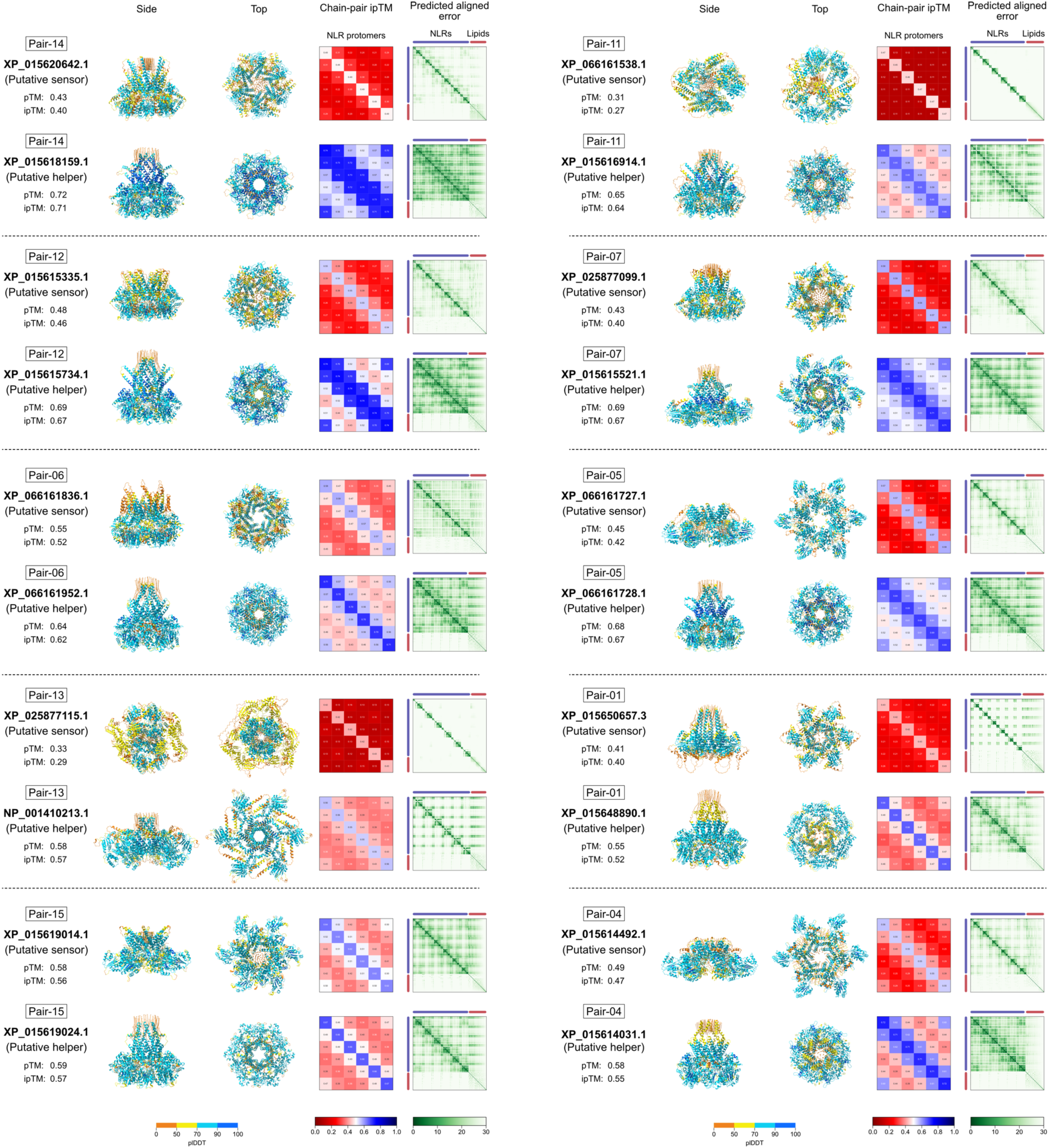
Hexameric AlphaFold 3 predictions for NLR pairs described by Stein et al., 2018. The amino acid sequences of the oligomerizing domains of the NLR proteins, from the N-terminus to the end of the NB-ARC domain, were used for the prediction. The hexameric structures were modelled with 50 oleic acids using seed value 1. The predicted structures were visualized with ChimeraX (Meng et al., 2023).

**Table S1. List of previously reported NLR pairs.** All sequences were derived from either RefPlantNLR (Kourelis et al., 2021) or NCBI. Sensor and helper NLRs are defined based on the presence or absence of an integrated domain, respectively. Domain architectures were annotated using NLRtracker (Kourelis et al., 2021). For domain architecture, “C,” “N,” “L,” and “O” represent the CC, NB-ARC, LRR, and other integrated domains, respectively. Note that InterProScan did not annotate an integrated domain of PIK5-NP as previously reported (Kourelis et al., 2021). Therefore, we manually replaced the domain architecture of PIK5-NP from “CNL” to “CONL” as it contains the HMA domain between the NB-ARC and LRR (Białas et al., 2021).

**Table S2. Summary of AlphaFold 3 predictions for previously reported NLR pairs.**

**Table S3. HMM scores of MADA motifs in previously reported NLR pairs.** The presence or absence of the MADA or MADA-like motif was analyzed using the HMM model in Adachi et al., 2019a. MADA, MADA-like, and no MADA motifs were classified based on HMM scores (> 10, < 10, and NA, respectively).

**Table S4. List of rice NLR pairs described by Stein et al., 2018.** The NLR pairs were identified from the rice cultivar Nipponbare genome annotation (Stein et al., 2018). Based on Supplementary Data 6 in Stein et al., 2018, the NLR pairs that i) are genetically linked in head-to-head orientations; ii) belong to distinct phylogenetic clades were extracted. Domain architectures were annotated using NLRtracker (Kourelis et al., 2021). Columns from Supplementary Data 6 in Stein et al., 2018 and those added in this study are highlighted in yellow and green, respectively.

**Table S5. BLASTP results using NLRs from Stein et al., 2018 as queries against those in the NCBI RefSeq annotation as subjects.** DIAMOND BLASTP (Buchfink et al., 2021) was used to identify corresponding sequences between the datasets. NLRs from Stein et al., 2018 and those in the NCBI RefSeq annotation of the rice cultivar Nipponbare (GCF_034140825.1) served as the query and subject sets, respectively.

**Table S6. Summary of NLRtracker outputs for rice NLR pairs from the NCBI RefSeq annotation of Nipponbare (GCF_034140825.1), corresponding to those described by Stein et al., 2018.** Only NLR pairs with a full N-terminal CC domain in both NLRs were processed for AlphaFold 3 predictions.

**Table S7. Summary of AlphaFold 3 predictions for rice NLR pairs from the NCBI RefSeq annotation of Nipponbare (GCF_034140825.1), corresponding to those described by Stein et al., 2018.** Putative sensors and helpers were assigned based on the average pTM scores of pentameric and hexameric structures, with a putative sensor having the lower average score and a putative helper having the higher average score.

**Data S1. Taxonomic and genome metadata of species used for phylogenetic analysis.**

**Data S2. Sequence and metadata of full-length and NB-ARC domains of NLR sequences used for phylogenetic analysis.**

**Data S3. Alignment of 5,509 NB-ARC sequences of NLRs used in Fig. S1.**

**Data S4. Phylogenetic tree of 5,509 NB-ARC sequences of NLRs used in Fig. S1.**

